# Identifying Causal Variants by Fine Mapping Across Multiple Studies

**DOI:** 10.1101/2020.01.15.908517

**Authors:** Nathan LaPierre, Kodi Taraszka, Helen Huang, Rosemary He, Farhad Hormozdiari, Eleazar Eskin

## Abstract

Increasingly large Genome-Wide Association Studies (GWAS) have yielded numerous variants associated with many complex traits, motivating the development of “fine mapping” methods to identify which of the associated variants are causal. Additionally, GWAS of the same trait for different populations are increasingly available, raising the possibility of refining fine mapping results further by leveraging different linkage disequilibrium (LD) structures across studies. Here, we introduce multiple study causal variants identification in associated regions (MsCAVIAR), a method that extends the popular CAVIAR fine mapping framework to a multiple study setting using a random effects model. MsCAVIAR only requires summary statistics and LD as input, accounts for uncertainty in association statistics using a multivariate normal model, allows for multiple causal variants at a locus, and explicitly models the possibility of different SNP effect sizes in different populations. In a trans-ethnic, trans-biobank Type 2 Diabetes analysis, we show that MsCAVIAR returns causal set sizes that are over 20% smaller than those given by current state of the art methods for trans-ethnic fine-mapping.

## Introduction

Genome-Wide Association Studies (GWAS) have successfully identified numerous genetic variants associated with a variety of complex traits in humans [1–3]. However, most of these associated variants are not causal, and are simply in Linkage Disequilibrium (LD) with the true causal variants. Identifying these causal variants is a crucial step towards understanding the genetic architecture of complex traits, but testing all associated variants at each locus using functional studies is cost-prohibitive. This problem is addressed by statistical “fine mapping” methods, which attempt to prioritize a small subset of variants for further testing while accounting for LD structure [4].

The classic approach to fine mapping involves simply selecting a given number of SNPs with the strongest association statistics for follow-up, but this performs sub-optimally because it does not account for LD structure [5]. Bayesian methods that did account for LD structure were developed [6, 7], but were based upon the simplifying assumption that each locus only harbors a single causal variant, which is not true in many cases [8]. Additionally, many early methods required individual-level genetic data, whereas many human GWAS often provide only summary statistics due to privacy concerns. CAVIAR [8] introduced a Bayesian approach that relied only on summary statistics and LD, accounted for uncertainty in association statistics using a multivariate normal (MVN) distribution, and allowed for the possibility of multiple causal SNPs at a locus. This approach was widely adopted and later made more efficient by CAVIARBF [9] and FINEMAP [10].

There is growing interest in improving fine-mapping by leveraging information from multiple studies. One of the most important examples of this is trans-ethnic fine mapping, which can significantly improve fine mapping power and resolution by leveraging the distinct LD structures in each population, as seen in methods such as trans-ethnic PAINTOR [11] and MR-MEGA [12]. Intuitively, the set of SNPs that are tightly correlated with the causal SNP(s) will be different in different populations, allowing more SNPs to be filtered out as potential candidates. However, the varying LD patterns also present a unique challenge in the multiple study setting that trans-ethnic fine mapping methods must handle. Additionally, while there is evidence that the same SNPs drive association signals across populations, there is also heterogeneity in their effect sizes, presenting another challenge [13]. Existing methods either assume a single causal SNP at each locus [12, 14] or do not explicitly model heterogeneity [11], limiting their power [15].

In this paper, we present MsCAVIAR, a novel method that addresses these challenges. We retain the Bayesian MVN framework of CAVIAR while introducing a novel approach to explicitly account for the heterogeneity of effect sizes between studies using a Random-Effects (RE) model. Our method requires only summary statistics and LD matrices as input, allows for multiple causal variants at a locus, and models uncertainty in association statistics and between-study heterogeneity. The output is a set of SNPs that, with a user-set confidence threshold (e.g. 95%), contains all causal SNPs at the locus.

We show in simulation studies that MsCAVIAR outperforms existing trans-ethnic fine mapping methods [11] and extensions of methods such as CAVIAR [8] to the multiple study setting. In a trans-ethnic, trans-biobank analysis of Type 2 Diabetes, we demonstrate that MsCAVIAR significantly improves the resolution of fine mapping compared to trans-ethnic PAINTOR or running CAVIAR on either population individually. MsCAVIAR is freely available at https://github.com/nlapier2/MsCAVIAR.

## Results

### MsCAVIAR overview

Our method, MsCAVIAR, takes as input the association statistics (e.g. Z-scores) for SNPs at the same locus in multiple studies and the linkage disequilibrium (LD) structure between variants obtained from in-sample genotyped data. MsCAVIAR computes and outputs a minimal-sized “causal set” of SNPs that, with probability at least *ρ*, contains *all* causal SNPs. This process is visualized in Figure 1.

**Fig. 1.**
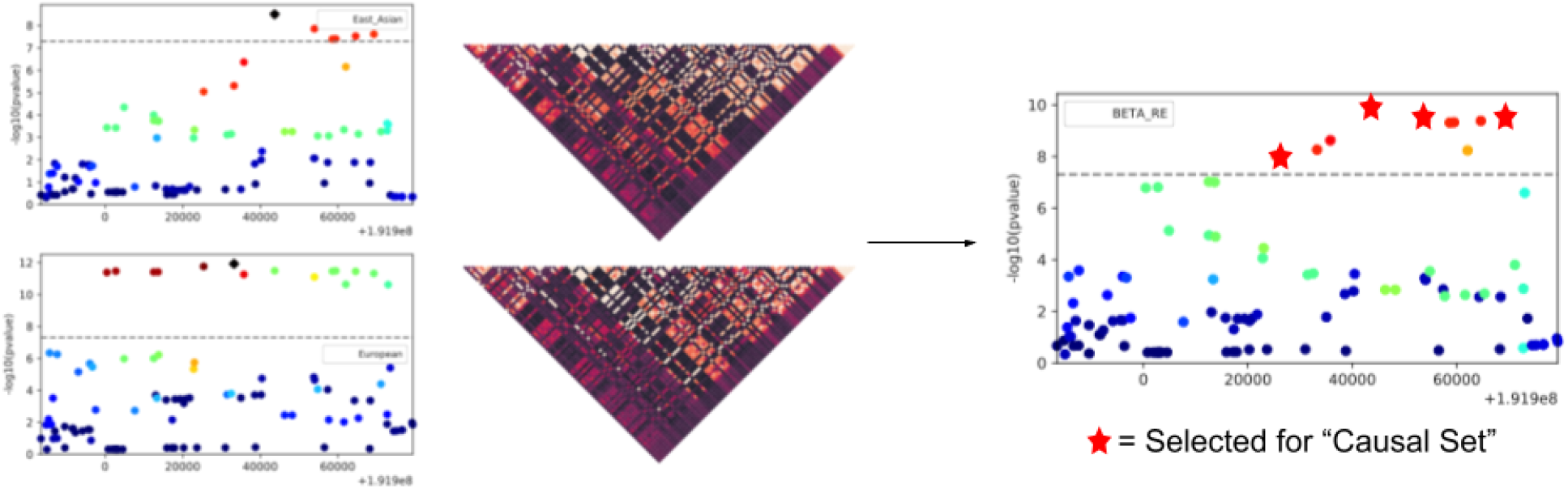
Overview of MsCAVIAR. MsCAVIAR takes as input the Z-scores and LD matrices for SNPs at a locus in two or more studies (left). Based on this input, MsCAVIAR leverages the different LD structures and models SNP effect heterogeneity to produce a refined “causal set” of SNPs (shown as red stars above) for follow-up functional validation studies. This set is often smaller than the set of SNPs that are significant in meta-analysis results (right).

By our definition of a causal set, every causal SNP must be contained in the set with high probability, but not every SNP in the set need to be causal. Concretely, each SNP can be assigned a binary causal status: 1 for causal or 0 for non-causal. So long as none of the SNPs outside of the causal set are set to 1, the assignments are compatible with our definition of a causal set. We can represent these causal status assignments in a binary vector with one entry for each SNP denoting its causal status; we call such a vector a “configuration” and denote it as *C*. For each configuration *C* compatible with the causal set, we compute its (posterior) probability in a Bayesian manner: the probability of a configuration of SNPs being causal given the association statistics can be computed by modeling a prior probability for that configuration and a likelihood function for the association statistics given the assumed causal SNPs given by *C* (see Methods for details).

The overall likelihood function can be decomposed into a product over the likelihood function for each study, since we assume that the studies are independent. More specifically, we assume that there is a true global effect size for a SNP over all possible populations, around which the effect sizes for that SNP in different studies are independently drawn according to a heterogeneity variance parameter (Methods). This allows MsCAVIAR to model the fact that effect sizes of a SNP across different studies are related, but not equal. Because we expect the summary statistics to be a function of their LD with the causal SNPs, the parameters of the likelihood function for each study are different, assuming the studies have different LD patterns. By computing the product over the likelihood of each study, we are able to account for their different LD patterns in determining the likelihood over all the studies.

The posterior probability for a causal set is then computed by summing the posterior probabilities of all compatible configurations, and then dividing by the sum of the posterior probabilities for all possible configurations. We start by assessing causal sets containing only one SNP, and continue increasing the size of the causal sets analyzed until one of them exceeds the posterior probability threshold *ρ*. In practice, *ρ* is set to a high value such as 95%.

### MsCAVIAR improves fine mapping resolution in a simulation study

In order to evaluate the performance of MsCAVIAR as compared with other methods, we performed a simulation study. In order to select realistic loci for fine-mapping, we identified regions in a trans-ethnic GWAS of rheumatoid arthritis [16] that contained peak SNPs with p-values of less than 0.0001 and contained ten or more SNPs in a 100kbp region centered around that peak. For each such locus, we used the 1000 Genomes project [17] to generate LD matrices for the SNPs at that locus for both European and East Asian populations. Out of these loci, we selected one region with relatively low LD, where 20% of the SNPs have LD equal to or higher than 0.5, and one region with relatively high LD, where 80% of the SNPs have LD equal to or higher than 0.5 (Figure 2, LD matrices). These represent easier and more difficult scenarios, respectively, for fine mapping, since LD makes signals more difficult to distinguish. We pruned groups of SNPs that were in perfect LD in one or more of the populations, leaving one SNP for each. If a group of SNPs were in perfect LD in one population, but not the other, we retained the SNP with the highest Z-score in the other population in order to retain the most signal.

**Fig. 2.**
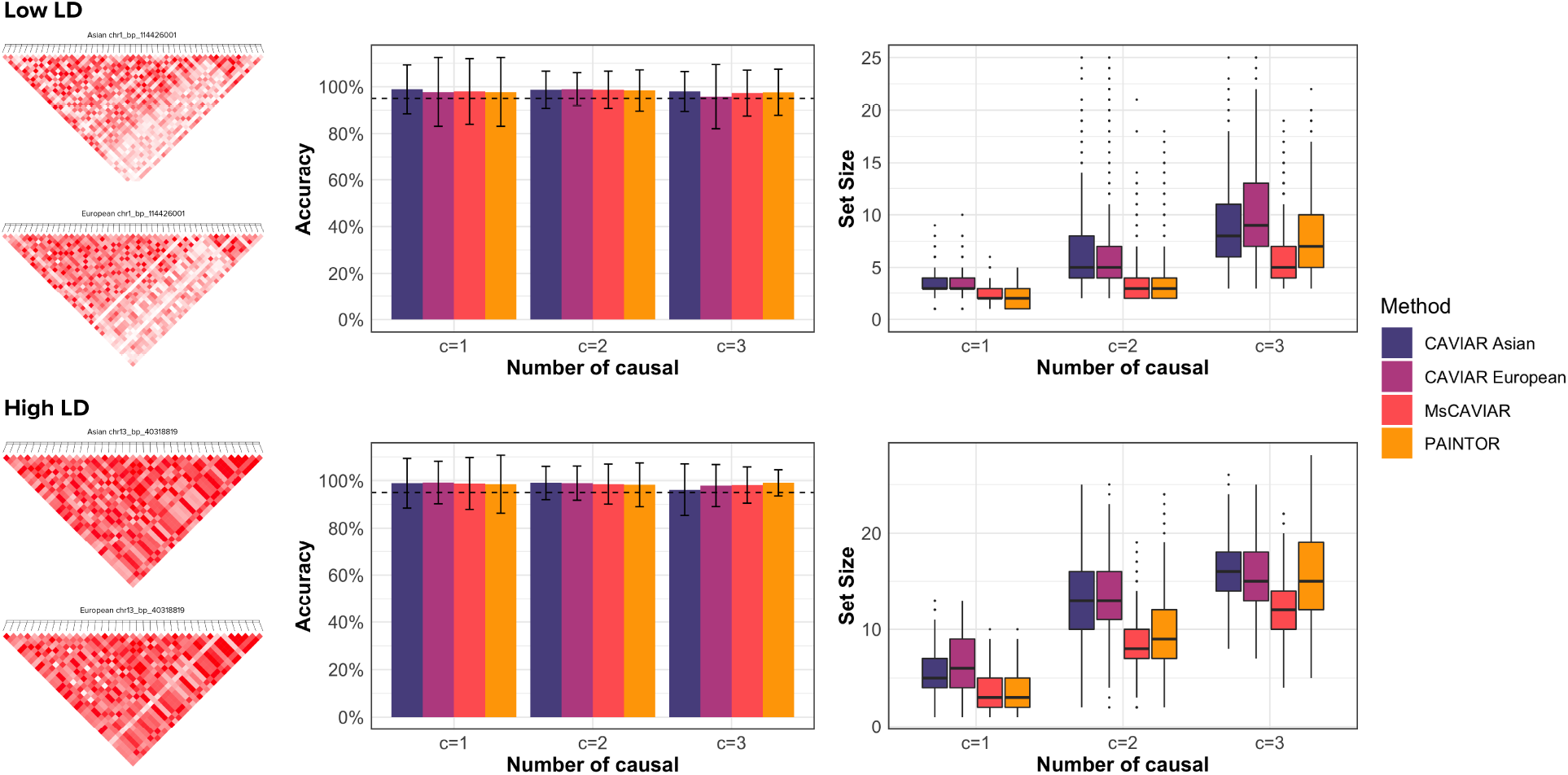
Comparison of accuracy and set size using simulated data. We simulated a trans-ethnic GWAS by using LD matrices generated from European and East Asian populations in the 1000 Genomes project. One relatively low LD region and one relatively high LD region were chosen (far left). Using these LD matrices, we implanted either 1, 2, or 3 causal SNPs and simulated their effect sizes. For each number of causal SNPs, we performed 1000 simulations (e.g. re-picking the causal SNPs and re-drawing the causal SNP effect sizes). In this figure, we report the average accuracy and standard deviation for each method in the bar graph (middle) and the set size in the box-plot (right). All methods were run with posterior probability threshold *ρ*^*∗*^ = 0.95, so methods with 95% or higher accuracy were considered “well-calibrated” (dashed line in the bar plots).

Using these LD matrices, we implanted causal SNPs and simulated their effect sizes. In each simulation, we implanted either 1, 2, or 3 causal SNPs. Each casual SNP’s true non-centrality parameter *Λ* was drawn according to 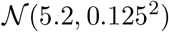. We then drew the non-centrality parameter for each study *i* according to 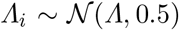, and subsequently the summary statistics according to 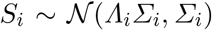. For each number of causal SNPs, we performed 1000 replicate simulations (e.g. re-drawing the causal SNP effect sizes and re-picking the causal SNPs).

Using this data, we compared MsCAVIAR to the trans-ethnic mode of PAINTOR [11] and to CAVIAR [8] run on East Asians and Europeans, individually (Figure 2). All methods were run with posterior probability threshold *ρ*^*∗*^ = 0.95, so methods with 95% or higher accuracy were considered “well-calibrated” (dashed line in the bar plots). MsCAVIAR’s heterogeneity parameter was set to *τ*^2^ = 0.5 (Methods); different settings for *τ*^2^ gave similar results. All methods were well-calibrated in both LD settings (Figure 2, bar plots). This is unsurprising since CAVIAR is well-known to be calibrated in the single study setting [8–10], as is PAINTOR in the trans-ethnic setting [11].

However, when considering the subset of simulations in which each method was able to correctly capture all causal variants (i.e. 100% accuracy), we observed that MsCAVIAR consistently returns the smallest average set size (Figure 2, box plots). MsCAVIAR and PAINTOR return smaller set sizes than CAVIAR run on either population across all settings, highlighting the value of using varying LD patterns in different populations to refine fine-mapping results. MsCAVIAR returned smaller set sizes than PAINTOR with multiple causal variants or high LD. This may be due to MsCAVIAR’s explicit modeling of heterogeneity between studies. In both the high LD and multiple causal variants setting, complex and strong correlations between non-causal and causal SNPs are induced, and modeling heterogeneity between studies allows for more effective use of the differing LD structures to disentagle non-causal from causal SNPs.

### MsCAVIAR is well-calibrated with different population sizes between studies

It is possible that input studies can have different sample sizes, in which case the effect sizes of their SNPs is expected to be different proportionally to sample size, in addition to heterogeneity. We tested whether MsCAVIAR would still be well-calibrated in this setting, and compared it again with trans-ethnic PAINTOR and with CAVIAR run on the individual populations (Figure 3).

**Fig. 3.**
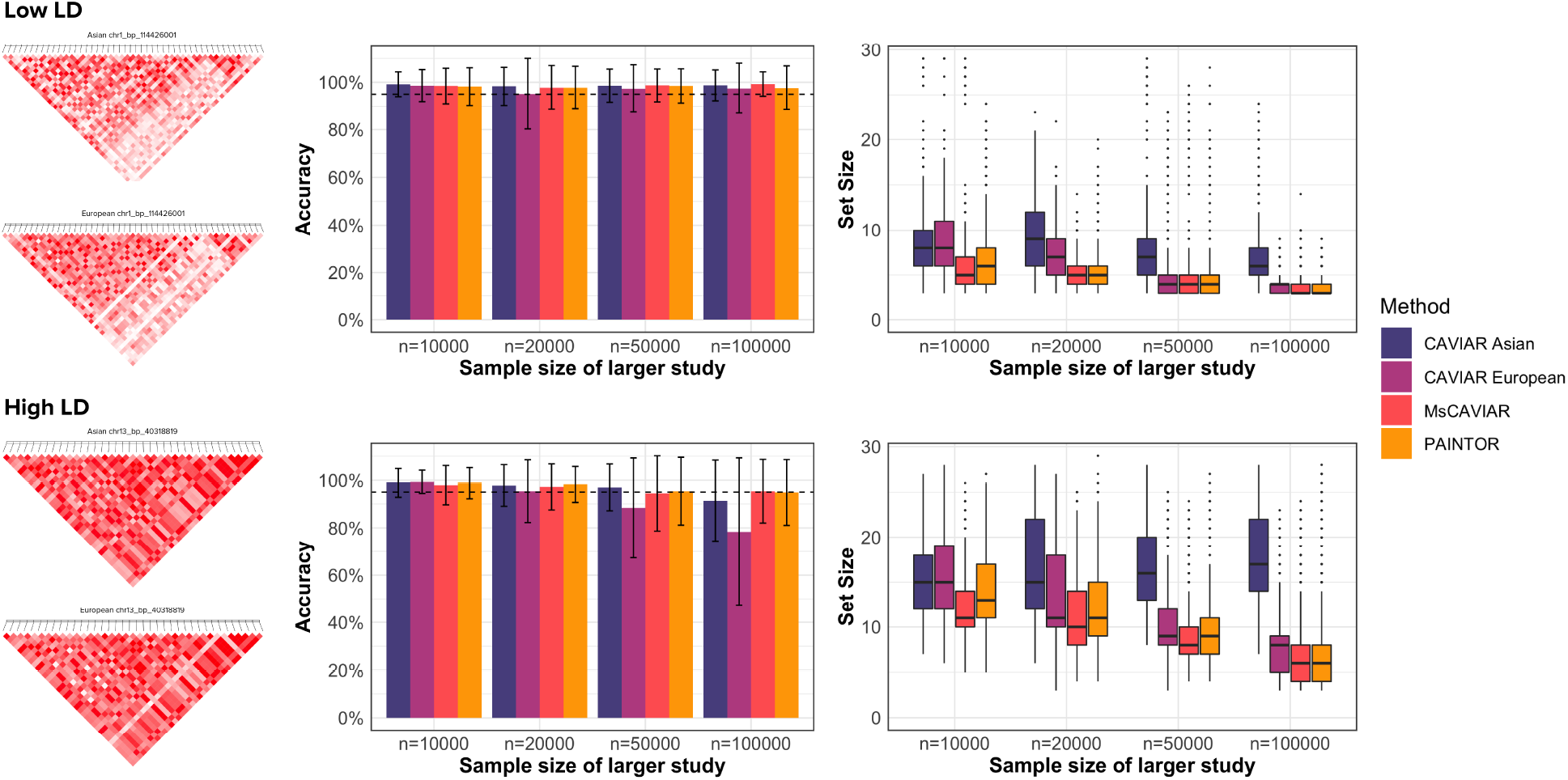
Comparison of accuracy and set size using simulated studies with unequal sample sizes. We simulated a trans-ethnic GWAS by using LD matrices generated from European and East Asian populations in the 1000 Genomes project. Using these LD matrices, we implanted 3 causal SNPs and simulated their effect sizes. In each set of simulations, we fix the population size of the Asian study at 10,000 individuals, and vary the European study to have population sizes of 1, 2, 5, or 10 times that of the Asian study. In this figure, we report the average accuracy and standard deviation for each method in the bar graph (left) and the set size in the box-plot (right).

In order to evaluate performance under this scenario in a simulation study, we used the same LD matrices from the previous section, but now varied the population size for one of the studies. We fixed the population size of the Asian study at 10,000 individuals, and varied the European study to have population sizes of 1, 2, 5, or 10 times that of the Asian study. Consequently, the effect sizes of causal SNPs in the European study were larger than those of the corresponding SNPs in the Asian study by a factor of 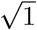, 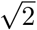, 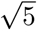 and 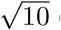 (Methods). For the sake of sufficient statistical power, we ensured that the causal variants in the smaller study were still statistically significant genome-wide. 1000 simulation replicates were run for each LD setting. In each simulation, we implanted three causal SNPs and simulated their effect sizes, with the association statistics of non-causal SNPs being based on their correlation with causal SNPs (Methods). All methods were run with posterior probability threshold *ρ*^*∗*^ = 0.95, so methods with 95% or higher accuracy were considered “well-calibrated” (dashed line in the bar plots). MsCAVIAR was run with its heterogeneity parameter set at *τ*^2^ = 0.5 (Methods).

Once again, MsCAVIAR was well-calibrated and generally returned the smallest causal set sizes. As the sample size difference grew, the difference between MsCAVIAR, CAVIAR on Europeans, and PAINTOR tended to diminish. This is likely due to the fact that we required SNPs to be genome-wide significant in the smaller study, such that the larger study had very large effect sizes for causal SNPs when there was a significant sample size imbalance, making the fine mapping problem easier. Reinforcing this interpretation is the fact that CAVIAR on Asians had consistently much larger causal set sizes than the other methods when the sample size imbalance was large.

All methods were well-calibrated in the low LD setting, but we observed that as the sample size increases with high LD that CAVIAR’s calibration on the larger population decreases. This is likely due to the extremity of the situation, with exceptionally large effect sizes in combination with the high LD setting.

### MsCAVIAR improves fine mapping resolution in trans-ethnic Type 2 Diabetes analysis

In order to evaluate the performance of MsCAVIAR on real data, we performed a trans-ethnic, trans-biobank fine mapping analysis of Type 2 Diabetes (T2D) using summary statistics from the UK Biobank (UKB) [18] and Biobank Japan (BBJ) [19] projects. These studies involved 361,194 and 191,764 people, respectively. Only White Europeans from the UK Biobank were used. To generate loci for fine mapping, we centered 100kbp windows around genome wide-significant peak SNPs (p-value ≤ 5 ∗ 10^*−*8^), discarding all SNPs with p-values above 0.0001, as they were highly unlikely to be informative. We applied several other filters to identify loci where trans-ethnic fine mapping was worthwhile. If a SNP was genome wide-significant in one ethnic GWAS but had a p-value above 0.0001 in the other ethnicity, we did not allow this to be a peak SNP, as information from the other ethnicity would be unlikely to help improve resolution in this case. In these instances, fine mapping within one population would be sufficient. We also excluded all loci with fewer than ten SNPs with p-values below 0.0001 in each study, as fine mapping is not as useful when there are few strongly associated SNPs. We excluded loci from chromosome six, where there were numerous statistically significant SNP effect sizes due to the presence of human leukocyte antigen (HLA) regions.

After these filters, five loci remained to be analyzed by trans-ethnic fine mapping. Linkage disequilibrium (LD) matrices were generated from the 1000 Genomes project [17], with “European” and “East Asian” as the population names, using the “CalcLD 1KG VCF.py” script from the PAINTOR [20] GitHub repository. As a final step, we pruned groups of SNPs that were perfectly correlated with each other in both studies (arbitrarily picking one SNP in the group to retain), as they provided identical information and would cause the LD matrix to be low-rank. The resulting five loci had 9, 10, 10, 42, and 42 SNPs.

We ran CAVIAR [8], the trans-ethnic mode of PAINTOR [11], and MsCAVIAR on these loci, and evaluated their causal set sizes, since these methods have been shown to be well-calibrated and no ground truth is available (Figure 4). For MsCAVIAR, we set the heterogeneity parameter *τ*^2^ (Methods) to 0.5. For CAVIAR, we evaluated its performence when applying it to only the Asian (BBJ) data or to only the European (UKB) data. For all methods, we set the posterior probability threshold *ρ*∗ to 80% and set the maximum number of causal SNPs to 3.

**Fig. 4.**
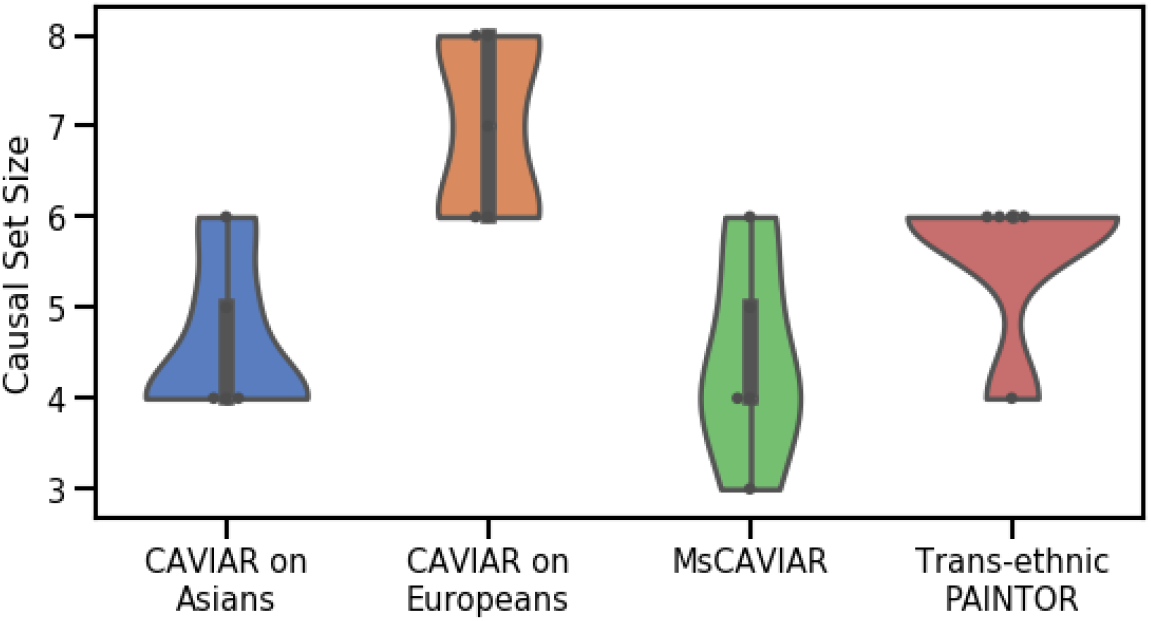
MsCAVIAR improves fine mapping resolution in trans-ethnic Type 2 Diabetes analysis. We compare the results of MsCAVIAR when applied to two Type 2 Diabetes (T2D) GWAS, White European people from the UK Biobank [18] and Japanese people from the Japan Biobank [19], versus trans-ethnic PAINTOR [11] and applying CAVIAR [8] to each population individually. The T2D data sets have five independent loci with at least one significant variant. The Y-axis is the size of the causal set for each locus. Each dot indicates the causal set for one locus. The violins represent the range of causal set sizes identified by each tool, and the width corresponds to the frequency of that causal set size for that tool.

Whereas the original five loci contained a combined 113 SNPs, MsCAVIAR yielded causal set sizes of 3, 4, 4, 5, and 6, for a total of 22 SNPs (80.53% reduction). CAVIAR on BBJ returned 23 SNPs, CAVIAR on UKB returned 35 SNPs, and Trans-ethnic PAINTOR returned 28 SNPs. Thus, MsCAVIAR yielded a reduction in combined set size of 4.34%, 37.14%, and 21.34%, respectively, compared to those alternative approaches. The Wilcoxon signed-rank test p-values for the alternate hypothesis that MsCAVIAR’s returned causal set sizes were smaller than those given by CAVIAR on BBJ, CAVIAR on UKB, and trans-ethnic PAINTOR were 0.159, 0.019, and 0.031, respectively. All methods ran in less than two minutes for all loci combined, with CAVIAR running the fastest, followed by MsCAVIAR, followed by trans-ethnic PAINTOR.

Based on these findings, the BBJ data seems to have an LD structure that is more favorable to fine mapping for these particular loci for T2D, since CAVIAR on BBJ refined loci much more than CAVIAR on UKB. However, MsCAVIAR was able to leverage both studies to achieve the best resolution. It is interesting that trans-ethnic PAINTOR refined loci less than CAVIAR on BBJ. This may be due to a lack of explicit modeling of heterogeneity and differing sample sizes between the studies in PAINTOR’s model. Overall, the finding that MsCAVIAR improves fine mapping resolution versus these other approaches is consistent with the findings in our simulation study.

## Discussion

In this work, we introduced MsCAVIAR, a method for identifying causal variants in associated regions while leveraging information from multiple studies. Our approach requires only summary statistics as opposed to genotype data and handles heterogeneity of effect sizes, differing sample sizes, and different LD structures between studies, making trans-ethnic fine mapping an ideal application. We demonstrated that our method is well-calibrated in simulation studies and improves fine-mapping resolution in both simulated and real data. MsCAVIAR is available as free and open source software (https://github.com/nlapier2/MsCAVIAR).

We make several important assumptions in this model, which may not always be true. While it has been shown that many causal SNPs are shared across populations [13], this may not always be the case. Ideally, this would be obvious from the summary statistics because the population in which the SNP is causal should have much more association signal, in which case one should just apply CAVIAR or a similar single-study method to that study. We also assume that all studies are drawn with equal heterogeneity *τ*^2^. This is unlikely to be true if multiple studies are from a single population while another study is from a different population. Since the primary benefit of MsCAVIAR is its utilization of varying LD structures, it is unlikely that multiple studies from the same population will confer much benefit. Therefore, we recommend using only one study from each population. However, it is still possible that even ostensibly different populations may be more similar to each other at certain loci than other populations. Therefore, we plan to extend our method to handle this case in future work.

## Methods

### Fine mapping in a single study

We now describe a standard approach for fine mapping significant variants from a genome-wide association study (GWAS). In the GWAS, let there be *n* individuals, all of whom have been genotyped at *m* variants. For each individual *j*, we measure a quantitative trait *y*_*j*_, resulting in the *n* × 1 column vector *Y* of phenotypic values. We denote *G* as the *n* × *m* matrix of the genotypes where *g*_*ij*_ ∈ {0, 1, 2} is the minor allele count for the *j*th individual at variant *i*. We standardize *G* according to the population proportion *p* of the minor allele and denote this as *X* where 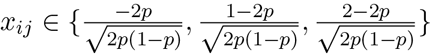.

We assume Fisher’s polygenic model, which means *Y* is normally distributed and each variant *x*_*i*_ has a linear effect on *Y*. We, therefore, have the following model:

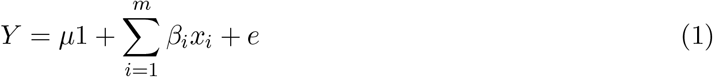

where *β*_*i*_ is the effect size variant *x*_*i*_ and *e* is variation in *Y* not explained by additive genetic effects. *e* is an *n* × 1 vector and follows the Gaussian distribution 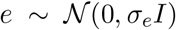 where each individual’s residual is independently and identically distributed. From this linear model, we define 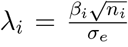, the standardized effect size of the *i*th variant, which is also referred to as the “non-centrality parameter” (NCP). Let *Λ* = [*λ*_1_… *λ*_m_] be an *m* × 1 column vector of non-centrality parameters. Furthermore, we define the the summary statistic *s*_*i*_ for each effect size *β*_*i*_, where 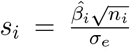, where 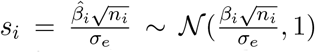. Let *S* be an *m* × 1 vector of summary statistics measured for each variant. As previously shown [8, 21], *S* follows a multivariate normal (MVN) distribution, 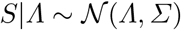 where *Σ* is the pairwise correlation structure between variants (LD). The expected value of each statistic *s*_*i*_ is a function of its correlation to a causal variant.

While the values of *Λ* are a function of a variant’s relationship to a causal variant, the vector itself does not indicate the variant’s causal status; therefore, we introduce an *m* × 1 binary vector *C* = {0, 1}^*m*^ for indicating whether a variant truly does have a non-zero effect size. We will now define *Λ*_*C*_ where each index 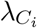 is as follows:

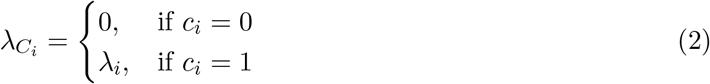

This means that if *C* = [1… 1], *Λ*_*C*_ = *Λ* because all variants would be causal. The distribution of *Λ*_*C*_ can be defined as:

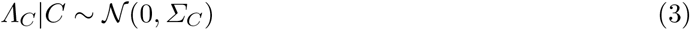

where

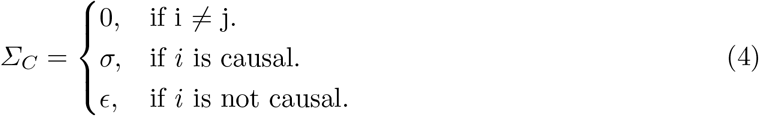

and where *ϵ* is a small constant to ensure that the matrix *Σ*_*C*_ is full rank. Here, and below, we use the shorthand *σ* to represent the variance of the 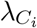 (see the subsection “Extending MsCAVIAR to different sample sizes” for details on this parameter). The off-diagonals of *Σ*_*C*_ are zero because the effect sizes of causal variants are independent of one another.

We will now more formally define *Λ* as *Λ* = *ΣΛ*_*C*_, which is to say the non-zero values in *Λ* are either due to the variant being truly causal or the variant’s correlation structure with the causal variant(s). This and the fact that LD structure is symmetric *Σ* = *Σ^T^* leads to the following distribution for *Λ*|*C*:

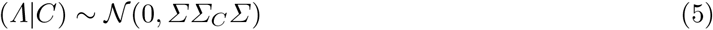

We will now define *γ* as the probability of a variant being causal, which makes the causal status for the *i*th variant a Bernoulli random variable with the following probability mass function: 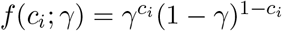. We assume the causal status for each variant is independent of the other variants, leading to the following prior for the our indicator vector: 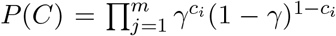. Assuming that each variant has a probability *γ* of having a causal effect, the prior can then be written as follows:

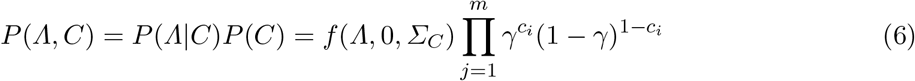

where *f* (*Λ,* 0*, Σ*_*C*_) is the probability density function shown in equation 5.

We determine which variants are causal by calculating the posterior probability of each configuration 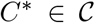, where 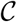 is the set of all possible configurations, given the set of summary statistics:

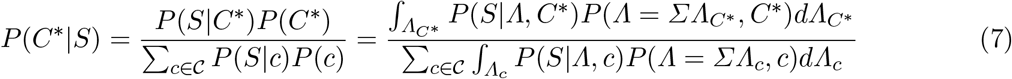

For us to calculate the posterior probability of *C^∗^* given *S*, we need to integrate over all possible values for the non-centrality parameters of the causal variants in *Λ* in order to get the values of *Λ* that makes observing *S* most probable.

Given a set of 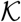 SNPs, 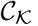, the posterior probability that this set of SNPs contains all the causal SNPs can then be calculated as follows:

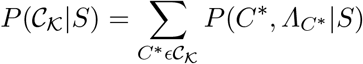

The goal is then to find the minimum-sized set 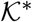 that has a posterior probability of at least *ρ*^*∗*^, called the “*ρ*^*∗*^ confidence set”:

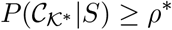

This is done by evaluating causal configuration vectors with only one non-zero element, and then those with two non-zero elements, and so on until the end condition above is met.

### Efficient computation of likelihood functions

The integral above is intractable. Fortunately, a closed-form solution is available due to the fact that, when a conjugate prior is multivariate normally distributed, its posterior predictive distribution is also multivariate normal. As shown above, 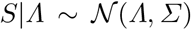 and 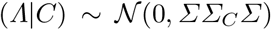. The posterior predictive form of *S* is then

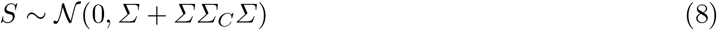

However, computing the likelihood of *S* with this distribution is still computationally expensive. Consider the multivariate normal probability density function, assuming the variable *Z* below is MVN distributed with mean *μ* and covariance matrix *Σ*:

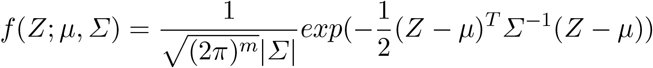

For *S*, the covariance matrix is *Σ* + *ΣΣ_C_ Σ*, which has dimension (*M* × *M*). Taking the determinant or inverse of this covariance matrix, as required by the above likelihood function, would take *O*(*m*^3^) time. Here, we demonstrate how to compute this likelihood efficiently, leveraging insights from several studies that have explored this topic [9, 10, 22].

We need to compute *S^T^* (*Σ* + *ΣΣ_C_ Σ*)^*−*1^*S* and |*Σ* + *ΣΣ_C_ Σ*| (note that our *μ* is 0). We can factor out *Σ* from both of the equations above:

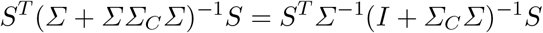

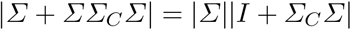

Notably, *S^T^ Σ^−^*^1^ and |*Σ*| can be computed once and re-used for every causal configuration *Σ*_*C*_. Below, we assume *Σ* is of full-rank; Lozano et. al [22] show how to address the low-rank case.

We use the Woodbury matrix identity [23], below, to speed up the matrix inversion equation:

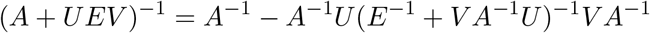

Here, we set *A* = *I_m×m_*, *E* = *I_k×k_*, and *UV* = *Σ_C_ Σ*. In particular, *U* is the (*m*×*k*) matrix of rows corresponding to causal SNPs in *Σ*_*C*_. We are taking advantage of the fact that rows corresponding to non-causal SNPs are zeros and thus do not affect the matrix multiplication. Similarly, *V* is the corresponding columns of *Σ*, and is (*k × m*). Applying the Woodbury matrix identity to our case, we get:

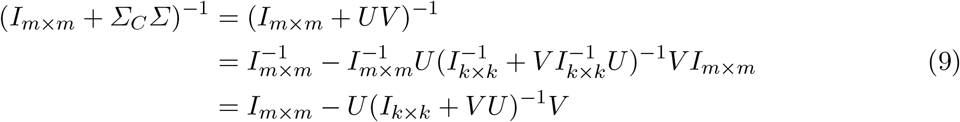

Crucially, we are now inverting a (*k* × *k*) matrix instead of an (*m* × *m*) matrix, where *k ≪ m* since most SNPs are not causal [22]. We use Sylvester’s determinant identity [24] to speed up the determinant computation as follows:

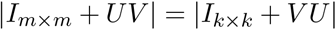

Similarly, we are computing the determinant of a (*k* × *k*) matrix instead of an (*m* × *m*) matrix. Using these speedups, the computation of the likelihood function of *S* is reduced from *O*(*m*^3^) to *O*(*k*^3^) plus some *O*(*mk*^2^) matrix multiplication operations, which is tractable under the reasonable assumption that each locus has at most *k* = 3 causal SNPs.

### Fine mapping across multiple studies

As GWAS continue to grow in size, frequency, and diversity, there is an increasing need for fine mapping methods that leverage results from multiple studies of the same trait. A simple approach is to assume that there is one true non-centrality parameter for every variant; therefore *Λ*_*C*_ is identical across studies. This approach is referred to as a fixed effects model. In this case, the *t*th study’s 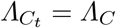.

While there is evidence that many causal SNPs are shared across populations [13], the assumption that the true causal non-centrality vector *Λ*_*C*_ is the same across studies is unrealistic, especially when the studies are measured in different ethnic groups [12, 14, 11].

We relax this assumption by utilizing a random effects model, in which each study *t* is allowed to have a different *Λ*_*Ct*_. Under this model, a causal SNP *q* has an overall mean non-centrality parameter, which we denote with the scalar *λ*_*q*_, from which the non-centrality parameter for SNP *q* in each study *t*, denoted by the scalar *λ*_*qt*_, is drawn with heterogeneity (variance) *τ*^2^. According to the polygenic model, *λ*_*q*_ is distributed as 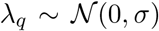; therefore, *λ*_*qt*_ is distributed as 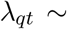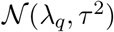. Consequently, the vector *Λ*_*Cq*_ for this SNP across all studies will have the following distribution:

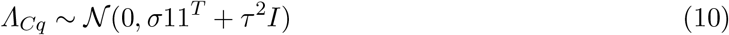

where *T* is the number of studies, 1 is a (*T* × *T*) matrix of 1s, and *I* is the (*T* × *T*) identity matrix. Intuitively, since the SNP *q* was drawn with variance *σ*, this variance component is shared across studies, while the variance component *τ*^2^ is study-specific and therefore it is only present along the diagonal of the covariance matrix. If a variant is not causal, its true effect size should be zero. We construct a matrix *Λ*_*C*_ of size (*mT* × *mT*), where *m* is the number of SNPs and each row corresponds to the *T*-length vector *Λ*_*Cq*_ corresponding to SNP *q*. In practice, we ensure that this matrix is full-rank by drawing the non-causal SNPs according to 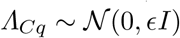, where *ϵ* is a small constant.

From this we will now build out the posterior probability of *P* (*C^∗^*|*S*) similarly to equation 7. Now instead of *Λ*_*t*_ = *Σ_t_Λ*_*C*_ for study *t*, we have to account for 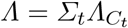 where *Λ*_*t*_ is drawn from a multivariate normal distribution. This means we have to integrate over the domain-space of 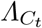 to as well as *Λ*_*C*_ to describe 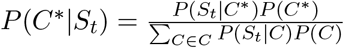

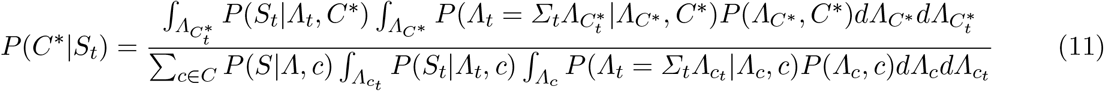

### Efficient meta-analysis

Now that we have described the distribution of each SNP in our meta-analysis, we show how to jointly analyze them. We begin by explicitly defining the structure of the covariance matrix between studies by way of a small example with three SNPs at a locus in two different studies. Since the covariance of a matrix is undefined, we denote *vec*(*Λ*_*C*_) as the vectorized form of the original matrix (*Λ*_*C*_). Concretely:

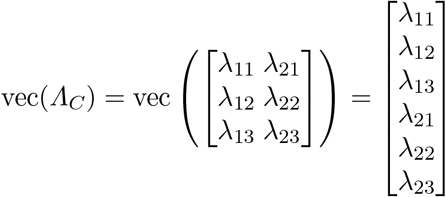

Assume SNPs 1 and 3 are causal and SNP 2 is not causal. Then the vectorized form of the non-centrality parameters given the causal statuses has the following multivariate normal distribution:

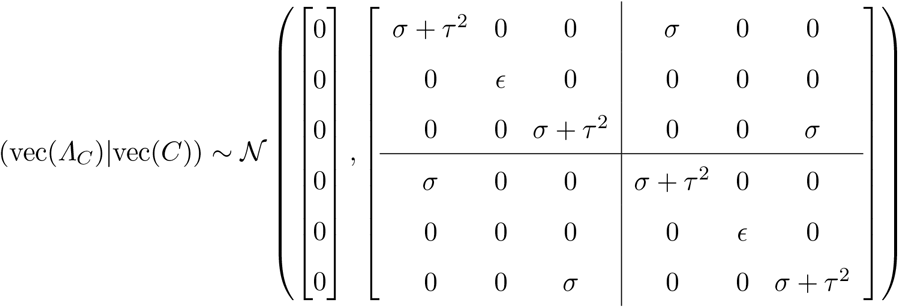

We call the covariance matrix above *Σ*_*C*_. Viewing *Σ*_*C*_ as having a block structure, the blocks along the diagonal represent SNPs from the same study, while off-diagonal blocks represent SNPs from different studies. Here *Σ*_*C*_ is (3 ∗ 2 × 3 ∗ 2) = (6 × 6); in general, for *m* SNPs and *n* studies, *Σ*_*C*_ will be (*mn* × *mn*). In other words, there will be an (*n* × *n*) grid of (*m* × *m*) blocks. Within each block, the diagonal represents each SNP’s variance, while the off-diagonal represents covariation between different SNPs. As SNPs are assumed to be independent, these are always 0. There are two variance components: the global genetic variance *σ* from which the global mean non-centrality parameter for a SNP is drawn, and the heterogeneity between studies *τ*^2^. When a SNP is causal, its variance (its covariance with itself in the same study) will contain both variance components (*τ*^2^ + *σ*), while its covariance with the same SNP in a different study will be *σ*, because they were drawn from the same overall non-centrality parameter with variance *σ* but were drawn separately with variance *τ*^2^.

The *Σ*_*C*_ above, leaving aside *c* for now, can alternately be written in the more-compact form

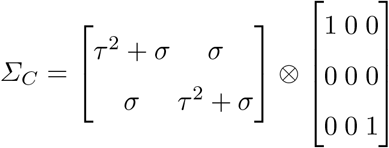

where ⊗ represents the Kronecker product operator. This can be further condensed and generalized into:

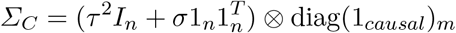

where *n* is the number of studies, *m* is the number of SNPs, 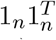 is the (*n* × *n*) matrix of all 1s, *I*_*n*_ is the (*n* × *n*) identity matrix, and *diag*(1_*causal*_)_*m*_ is an (*m* × *m*) diagonal matrix whose diagonal entries are given by the (1 × *m*) indicator vector 1_*causal*_ whose entries *i* are 1 if SNP *i* is causal and 0 otherwise.

As with CAVIAR, the *c* entries along the diagonal are small numbers to ensure full rank. Also note that the CAVIAR model is a specific case of this model, in which there is only one study and thus there is no *τ*^2^ component. The CAVIAR *Σ*_*C*_ has the same structure as the upper left block in the *Σ*_*C*_ above, when there are 3 SNPs and *τ*^2^ is set to 0.

The efficient computation properties for the single-study case also apply to the multiple-study case. In the latter setting, the matrices that need to be inverted are (*mn* × *mn*) instead of (*m* × *m*), where *m* and *n* are the number of SNPs in a locus and the number of studies, respectively. Consequently, in the Woodbury matrix identity equations, *U* and *V* are (*mn* × *kn*) and (*kn* × *mn*), respectively, where *k ≪ m* is the number of causal SNPs, and the matrix given by the Woodbury identity is (*kn* × *kn*). Sylvester’s determinant identity gives a matrix of this size as well. The computation time is thus reduced from (*mn* × *mn*) to (*kn* × *kn*).

### Extending MsCAVIAR to different sample sizes

Previously, we have assumed the *i*th SNP has one true mean non-centrality parameter, *λ*_*i*_. This simplifying assumption implies that all studies have the same sample size *n*, as the non-centrality parameter is a function of sample size 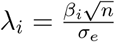. We are able to relax this assumption in MsCAVIAR to accommodate studies with differing sample sizes and in doing so will need to more precisely define *σ*. Additionally, we can no longer assume the summary statistic 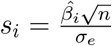 is drawn according to the global mean *λ*_*i*_; instead we will need to assume that study *m*’s observed effect size 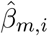 is an unbiased estimate of the true effect size of the SNP, *β*_*i*_, across studies where

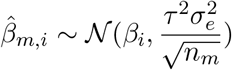

We now draw the true effect size of the *i*th SNP according to 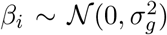 therefore, the mean effect size is 0 and the variance in effect size is the variance explained by this variant under an additive model, which we denote with 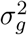. We will again draw the *m*th study’s non-centrality parameter for variant *i* according to this model. Each study *m* has its own sample size *n*_*m*_, environmental component 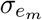, and we draw it with heterogeneity parameter *τ*^2^ as previously defined, so

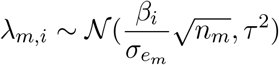

We will now operate under the standard assumption that *σ*_*e*_ has been standardized (*σ*_*e*_ = 1), so

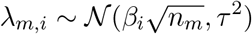

Using our previous definition for a single study, we now have

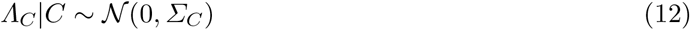

where

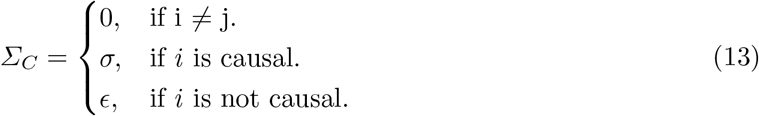

We now define *σ* more formally to be *σ_g_n*_*m*_ for the *m*th study. When we consider our matrix

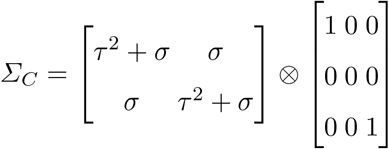

The *σ* along the diagonal is defined identically to the precise single study definition; however, when modeling multiple studies, this adjustment changes the covariance between causal variant for two studies. We now define 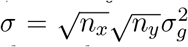 for two studies *x* and *y* with population sizes *n*_*x*_ and *n*_*y*_. Note that if two studies have the same population size *n*, we get the original definition of 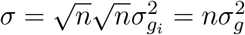 (recalling that here we are assuming 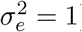.

### Parameter setting in practice

Traditionally, the effect size 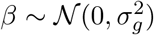 would be derived as a notion of the per-snp heritability. Here we do not define 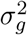 as such, but rather treat it as an abstraction: we avoid making any assumptions on how heritable the given trait is and how that heritability is partitioned between loci. The way we set this parameter in practice is as a parameter for statistical power. If study *m* has the smallest sample size, we set this value such that 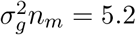 for all variants. Then the NCP for variant *i* in the corresponding study *m* is 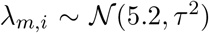. For another study *x* with larger sample size, its NCP is drawn as 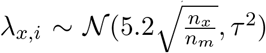. This value of 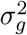 may not represent the actual heritability partitioning, but we set the parameter this way in our method for the practical purpose of giving MsCAVIAR power to fine map borderline significant variants in the smallest study.

